# Analysis of organelle content supports a membrane budding model of platelet biogenesis

**DOI:** 10.1101/2022.05.21.492939

**Authors:** Antoine F. Terreaux, Alison M. Farley, Christine Biben, Oliva Stonehouse, Samir Taoudi

## Abstract

Understanding how *in vivo* platelet biogenesis is undertaken is critical to making on-demand platelet production for clinical use feasible. We recently described the discovery of plasma membrane budding as a major *in vivo* platelet-producing pathway. *In vitro* recapitulation of this finding could pave the way towards efficient laboratory-based platelet production.

The plausibility of the plasma membrane budding model has been called into question. The foundation of this is the contention that the size and payload composition of plasma membrane buds are not consistent with *bona fide* platelets. Thus, membrane buds likely represent stages in megakaryocyte-derived microparticle formation.

Using 3D super-resolution imaging, we have performed a quantitative comparison of size and organelle content of plasma membrane buds, platelets, and microparticles in the adult mouse bone marrow. We unequivocally demonstrate that the structures we previously described as membrane buds exhibit the same size range as free platelets, that all buds contain organelles, and that membrane buds and free platelets contained an equivalent number of organelles. Crucially, membrane buds and microparticles are completely distinct from each other. To prevent future confusion between the processes of microparticle formation and platelet biogenesis, we propose using the more specific term “pre-platelet membrane buds”.

## Introduction

The current generations of platelet producing technologies are designed to recapitulate the process of proplatelet formation - the extension of long pseudopodia processes from megakaryocytes. These platforms can generate at 70 – 80 platelets per megakaryocyte ^1-3^. However, the *in vivo* platelet biomass is thought to be sustained by the production of approximately 1,000 – 3,000 platelets per megakaryocyte ^4,5^. This indicates that substantial additional refinement of *in vitro* proplatelet formation is required, or that other mechanisms drive the high rate of *in vivo* platelet production.

Using quantitative 3D imaging, 4D intra-vital imaging, and the use of a genetic model of severe thrombocytopenia, we proposed that the *in vivo* platelet biomass cannot be explained by proplatelet formation alone ^5^. We identified platelet production via plasma membrane budding (in which platelet-sized structures are produced and released directly from the megakaryocyte plasma membrane) as a mathematically adequate explanation for the physiological maintenance of the platelet biomass. We further argued that the platelet-like patterning of key cytoskeleton proteins (F-ACTIN, NMIIA, and TUBULIN) indicated that membrane buds were poised to be released as structurally competent platelets.

Recently, the plausibility of platelet production via membrane budding has been called into question. Based on the analysis of ultra-thin sections of adult bone marrow by transmission electron microscopy, Italiano *et al* ^6^ raised important criticisms of our study. The authors argued that the allotment of organelles (specifically α-granules) in platelets were not present in membrane buds. Additionally, the authors postulated that membrane budding was an indiscriminate process that did not yield products of size comparable to the uniformity of platelets. Italiano *et al* concluded that membrane budding could not represent a developmental stage in platelet biogenesis but were associated with microparticle (also known as microvesicle) formation.

Megakaryocyte/platelet-derived microparticles are 100 – 1,000 nm bodies ^7,8^ that can contain organelles such α-granules and mitochondria ^9^. As alluded to by Italiano *et al* ^6^, based on the difference in size between platelets and microparticles alone, it would be expected that microparticles contain far fewer organelles than platelets. Intriguingly, microparticle formation from megakaryocytes appears to involve plasma membrane budding ^7^.

Using full-volume super-resolution images of platelets, microparticles, and plasma membrane buds in the adult bone marrow, we compared size and organelle content and found that the membrane bud structures that we previously described as platelet precursors ^5^ are not consistent with microparticles but are statistically indistinguishable from *bona fide* platelets. These data provide critical evidence that supports the conclusion that plasma membrane budding is a major pathway of *in vivo* platelet biogenesis.

## Methods

### Mice

8 – 12-week-old C57BL/6 mice were used. Procedures were approved by the WEHI Animal Ethics Committee.

### Imaging

Femurs were prepared as previously described ^5^. Following cryopreservation in 30% sucrose, samples were snap frozen in OCT compound. 10 μm longitudinal sections were cut. Staining was performed for 16 hrs with either anti-GPB1α (R300, Emfret Analytics) and - VWF (A0082, Dako) antibodies, or anti-GPB1α and -Tom20 (sc11415, Santa Cruz) antibodies. Data were collected using the Airyscan function of a Zeiss LSM 980 confocal microscope (x63 objective lens, NA 1.4). Metrics were collected with the aid of Imaris and FIJI ^10^ software.

### Statistical analysis

*“n”* designates the number of independent experiments. “*o”* designates the number of observations. Graph production and analysis was performed with Prism.

## Results and Discussion

We previously described plasma membrane budding as the formation of platelet precursors at the surface of megakaryocytes that are released without proplatelet formation or cell death ^5^. An essential feature of this definition was bud size relative to free platelets (Figure 1A and Movie S1). To distinguish platelet production from other processes that might involve budding, we will refer to these structures as pre-platelet membrane buds.

**Figure 1.**
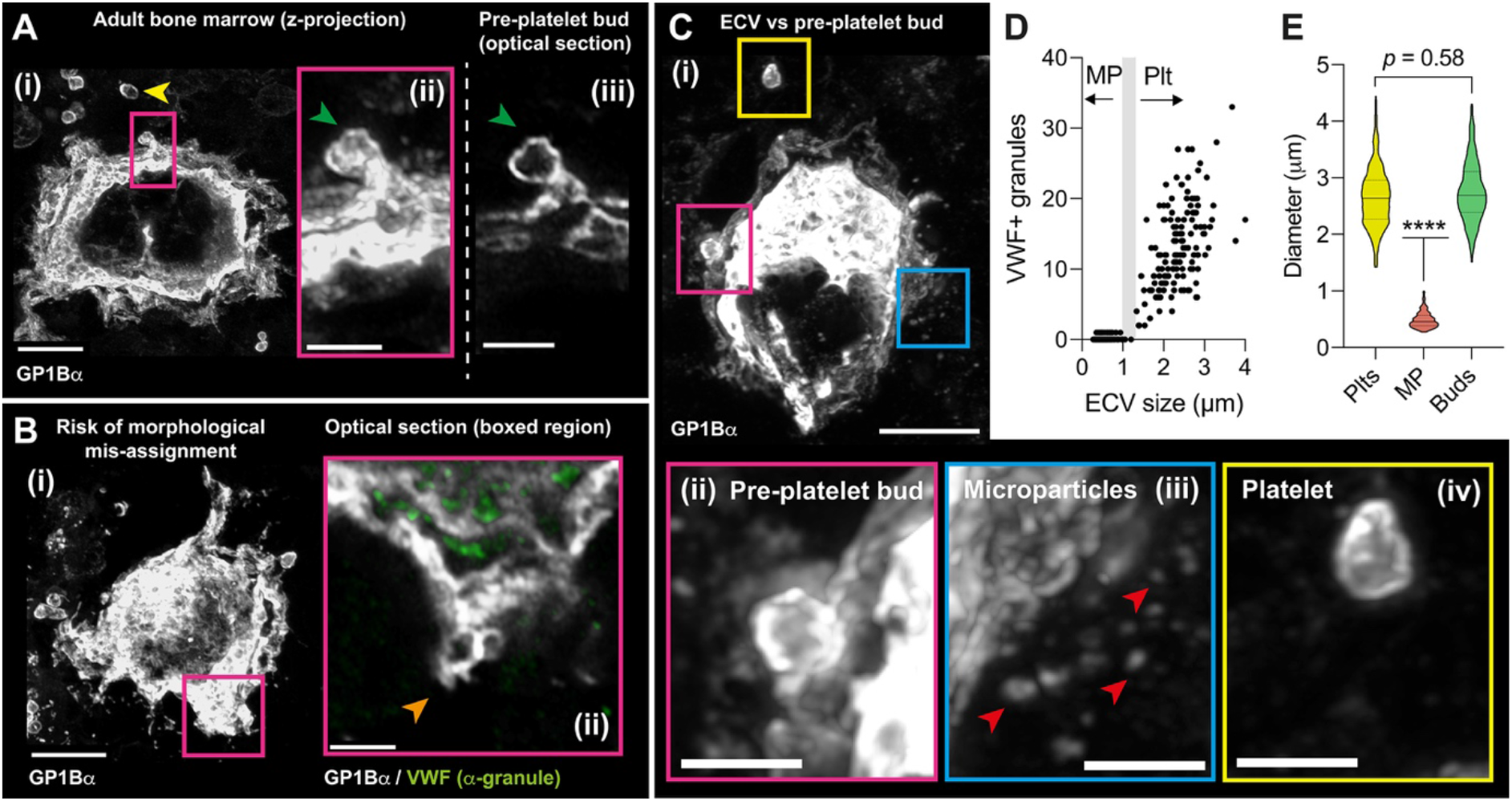
*In situ* distinction between pre-platelet membrane buds and megakaryocyte/platelet-derived microparticles in the adult bone marrow. **(A) (i)** 3D z-projection of a representative pre-platelet plasma membrane bud (boxed region). Note the comparable size of the pre-platelet bud to free platelets (yellow arrowhead). Scale bar, 10 μm. **(ii)** Enlargement of boxed region in **(i)**. Scale bar, 3 μm. **(iii)** Connection of the platelet-size membrane bud to the parental megakaryocyte is confirm by the study of optical sections. Scale bar, 3 μm. Data is representative of 99 buds from 5 independent adult mice. Anti-GP1Bα staining in grey. **(B)** Full-volume 3D context is critical to accurately identify pre-platelet plasma membrane buds. **(i)** Representative example of a non-budding bone marrow megakaryocyte. Scale bar, 10 μm. **(ii)** Single optical sections may appear to contain bud-like morphology (orange arrowhead). Scale bar, 3 μm. However, this interpretation does not hold when 3D context is revealed (as seen in **(i)** and Movie S1). As informed by *in vitro* and *in vivo* imaging^3^, *bona fide* pre-platelet membrane buds are recognised as platelet-size structures in continuity with the parental megakaryocyte (as in Fig. 1A). Anti-VWF staining shown in green, anti-GP1Bα in grey. **(C)** Pre-platelet membrane buds and microparticles are unequivocally distinguished according to size. **(i)** Budding megakaryocyte surrounded by microparticle-sized objects and a platelet. Scale bar, 10 μm. **(ii)** Enlargement of magenta boxed region in **(i)** of a pre-platelet membrane bud. Scale bar, 3 μm. **(iii)** Enlargement of blue boxed region in **(i)** of megakaryocyte/platelet microparticles. Scale bar, 3 μm. **(iv)** Enlargement of yellow boxed region in **(i)** of a free platelet. Scale bar, 3 μm. **(D)** Scatter plot of GP1Bα+ extra-cellular vesicle (EV) sizes and VWF+ α-granule content in sections of adult bone marrow. EV ≤ 1 μm were considered as microparticles (MP). EV ≥ 1.3 μm were considered as platelets (Plt); these always contained VWF+ α-granules. Grey bar represents EVs between 1 – 1.3 μm, these were excluded from analysis. (*o* = 340; *n* = 5 independent mice) **(E)** Violin plot of size comparison between free platelets, megakaryocyte/platelet-derived microparticles, and pre-platelet membrane buds in the adult bone marrow. (*o* = plts: 420, MPs: 188, buds: 125; *n* = 5 independent mice). Data were analysed using the Kruskal-Wallis test and Dunn’s multiple comparisons test. ****, *p* <0.0001. For each plot, the solid horizontal line = median, dashed lines = quartiles.

A critical step in the identification of pre-platelet buds is the acquisition of full-volume data to allow complete 3D reconstructions. 3D data is essential to prevent the mis-assignment of pre-platelet membrane bud identity to proplatelet extensions or spurious structures at the plasma membrane which are devoid of von Willebrand factor (VWF)-containing α-granules and do not conform to the classification of a pre-platelet bud (Figure 1B and Movie S2). Key to this is the presentation of a spheroid platelet-sized swelling attached to the plasma membrane of a megakaryocyte (Figure 1A). To make this assessment, we first calibrated expectation of size based on the free platelets captured in the tissue section. We have found that the use of physical ultra-thin sections (such as those used by Italiano *et al* ^6^) to be inadequate for the study of *in vivo* pre-platelet buds because they do not provide sufficient 3D context.

We generated full-volume 3D images of wildtype mouse bone marrow using Airyscan super-resolution microscopy. This modality provides a spatial resolution of 140 nm in *x* and *y* dimensions and 400 nm in *z* (at 488 nm excitation). This provided sufficient spatial resolution to identify VWF+ α-granules and Tom20+ mitochondria ^11-13^ in free platelets, 300 nm – 1000 nm microparticle-like structures ^7,9,14^, and pre-platelet buds ^5^ (Figure 1C).

To investigate the concern that the size range of pre-platelet buds is incompatible with being a platelet precursor, we first compared the size of platelets, microparticles, and pre-platelet buds. To establish microparticle and platelet benchmark values we quantified the maximal diameter and α-granule content of full-volume GP1Bα+ extra-cellular vesicles (Figure 1D). Due to technical limitations, we imposed a lower limit threshold of 300 nm, Megakaryocyte/platelet-derived microparticles are defined as structures < 1 μm in diameter ^7,9,14^. As expected ^9,14^, microparticles infrequently contained granules (Figure 1D). We based the threshold for platelet classification on the minimum diameter after which all extra-cellular vesicles contained granules; this was set at > 1.3 μm (Figure 1D). Microparticles (median diameter 0.5 μm, range 0.3 – 1.0 μm) were significantly smaller than platelets (median diameter 2.6 μm, range 1.4 – 4.4 μm) and pre-platelet buds (median diameter 2.7 μm, range 1.5 – 4.3 μm). The difference in size between free platelets and pre-platelet buds was not statistically significant (Figure 1E). This is consistent with pre-platelet buds being a direct ancestor to platelets but not to microparticles.

Using 3D electron microscopy, the absolute number of α-granules and mitochondria in mouse platelets have been determined ^11,15^. Using these data as a benchmark, we used Airyscan super-resolution confocal microscopy to accurately quantify α-granules and mitochondria content (Figure 2A - E). Given the limitations inherent to light microscopy in separating structures close together, it is possible that our data might underestimate the total number of organelles.

**Figure 2.**
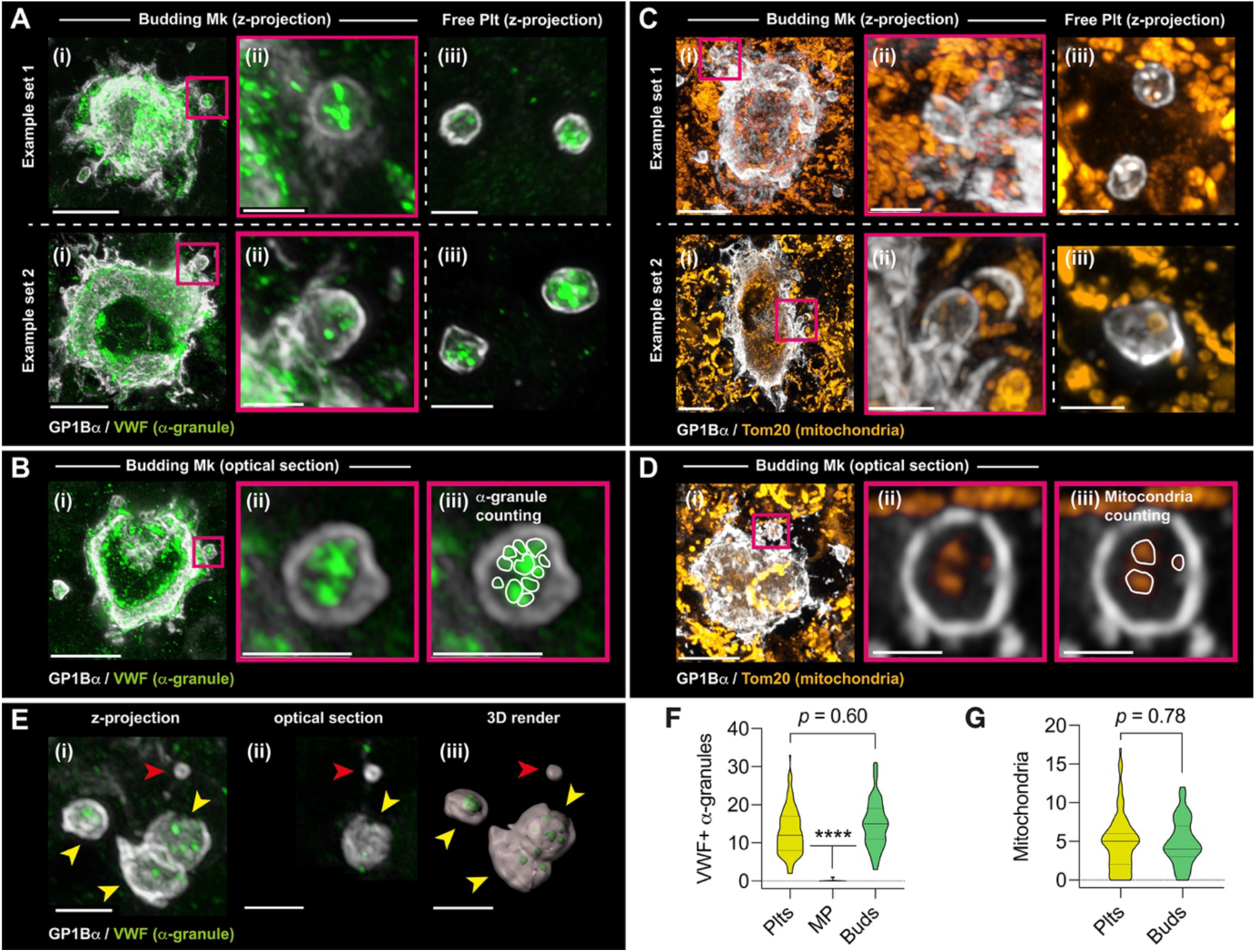
Pre-platelet membrane buds contain platelet-like numbers of α-granules and mitochondria. **(A)** Representative examples demonstrating the range of VWF+ α-granule distribution in pre-platelet membrane buds (i and ii) and free platelets (iii). Shown in (i) are examples of megakaryocytes with pre-platelet buds. (ii) Zoomed regions of pre-platelet buds from the boxed regions in (i). (iii) Free platelets present in the same field of view in megakaryocyte in (i). Grey, GP1Bα; Green, VWF. Scale bars for (i), 10 μm; scale bars for (ii) and (iii), 3 μm. **(B)** Representative example illustrating how VWF+ α-granules were quantified in pre-platelet membrane buds and free platelets. Grey, GP1Bα; Green, VWF. (i) Representative example of a megakaryocyte with pre-platelet bud. (ii) Zoomed region of pre-platelet buds from the boxed regions in (i). (iii) Superimposition of boundary lines (dashed) to highlight identified VWF+ granules. Grey, GP1Bα; Green, VWF. Scale bars for (i), 10 μm; scale bars for (ii) and (iii), 3 μm. **(C)** Representative examples demonstrating the range of mitochondria (anti-Tom-20 stained) distribution in pre-platelet membrane buds and free platelets. Shown in (i) are examples of megakaryocytes with pre-platelet buds. (ii) Zoomed regions of pre-platelet buds from the boxed regions in (i). (iii) Free platelets present in the same field of view in megakaryocyte in (i). Grey, GP1Bα; Orange, Tom20. Scale bars for (i), 10 μm; scale bars for (ii) and (iii), 3 μm. **(D)** Representative example illustrating how Tom-20+ mitochondria were quantified in pre-platelet membrane buds and free platelets. (i) Representative example of a megakaryocyte with pre-platelet bud. (ii) Zoomed region of pre-platelet buds from the boxed regions in (i). (iii) Superimposition of boundary lines (dashed) to highlight identified Tom20+ mitochondria. Grey, GP1Bα; Orange, Tom20. Scale bars for (i), 10 μm; scale bars for (ii) and (iii), 3 μm. **(E)** Representative example of VWF+ α-granule distribution in free platelets and microparticles in the bone marrow. (i) Representative example of VWF+ α-granules content in free platelets (yellow arrows) and microparticles (red arrow). (ii) Optical section of image in (i). (iii) 3D render of data in (i). Grey, GP1Bα; Green, VWF. Scale bars, 3 μm. **(F)** Violin plots of VWF+ α-granules in bone marrow free platelets (Plts), microparticles (MP), and pre-platelet membrane buds. *o* = 64 Plts, 65 MP, and 68 buds. *n* = 5 independent mice. Data were analysed using the Kruskal-Wallis test and Dunn’s multiple comparisons test. ****, p <0.0001. For each plot, the solid horizontal line = median, dashed lines = quartiles. **(G)** Violin plots of Tom20+ mitochondria in free platelets (Plts), microparticles (MP), and pre-platelet membrane buds. The high density of Tom20+ mitochondria in section prohibited accurate quantification of mitochondria in MPs. *o* = 50 Plts, 57 MP, and 54 buds. *n* = 5 independent mice. Data were analysed using the Mann-Whitney test. ****, p <0.0001. For each plot, the solid horizontal line = median, dashed lines = quartiles.

Platelets contained approximately 12 VWF+ α-granules (coefficient of variation [COV], 45%) and 5 mitochondria (COV, 73%) (Figure 2 F and G). This was in keeping previously described benchmark values ^11,15^. In contrast, microparticles rarely contained VWF+ granules (median of 0) (Figure 2 E - G). Due to the density of mitochondria in the bone marrow, mitochondria could not be accurately quantified in microparticles. Pre-platelet buds contained 15 α-granules (COV, 41%) and 4 mitochondria (COV, 60%), this was similar to the organelle content of free platelets (Figure 2 F and G).

These data unequivocally demonstrate that pre-platelet buds cannot be confused with microparticles and that they are comparable in size and organelle content to free platelets. These findings further support that membrane budding is a major mechanism of *in vivo* platelet biogenesis.

## Acknowledgments

We thank Douglas J Hilton for comments on the manuscript. This work was supported by an NH&MRC project grant (GNT2011770), a Speedy Innovation Grant, and Independent Research Institutes Infrastructure Support Scheme grant 361646 from the NH&MRC, the Australian Cancer Research Fund, and Victorian State Government Operational Infrastructure Support. ST was supported by a Fellowship from the Lorenzo and Pamela Galli Charitable Trust.

## Author contributions

AT and AF designed experiments and analyzed data. AT performed experiments. CB and OS provided essential input into experimental design and data analysis. ST conceived the study, designed experiments, and analyzed data. All authors contributed to manuscript preparation.

